# Invariant Odor Recognition with ON-OFF Neural Ensembles

**DOI:** 10.1101/2020.11.07.372870

**Authors:** Srinath Nizampatnam, Lijun Zhang, Rishabh Chandak, Nalin Katta, Barani Raman

## Abstract

Invariant recognition of a stimulus is a challenging pattern-recognition problem that must be dealt with by all sensory systems. Since neural responses evoked by a stimulus could be perturbed in a multitude of ways, could a single scheme be devised to achieve this computational capability? We examined this issue in locust olfactory system. We found that odor-evoked responses in individual projection neurons in the locust antennal lobe varied unpredictably with repetition, stimulus dynamics, stimulus history, presence of background odorants, and changes in ambient conditions. Yet, a highly-constrained Bayesian logistic regression approach with ternary weights could provide robust odor recognition. We found that this approach could be further simplified: sum firing rates of ON neurons and subtract total activity in OFF neurons (‘ON minus OFF’ classifier). Notably, we found that this approach could be generalized to develop a Boolean neural network that can perform well in a non-olfactory pattern recognition task.

## INTRODUCTION

Robustly recognizing a sensory stimulus is a necessity for the survival and propagation of all animals. Since this capability is demonstrated in all sensory systems, this raises the following question: what is the neural basis that underlies this feat of pattern recognition? Most stimuli are encountered in a multitude of ways in natural environments. Often, stimulus features such as intensity, duration, and recurrence could vary. In addition, external perturbances due to changes in environmental conditions (such as changes in humidity or temperature), the presence of other competing cues, or the temporal context (i.e. when it is received in a stimulus sequence) could also change independent of the variation in stimulus-specific features. An additional degree of interference can arise from changes in the sensory circuit due to plastic changes arising either from prior exposures or co-occurrence with other sensory cues. Given the complexity in carrying out the basic task of recognizing a stimulus, we wondered if there exists a computational framework that can compensate for all these disparate sources of variation and allow robust recognition of a stimulus. In particular, we sought to examine this issue in the well-studied locust olfactory system [1-13].

In the locust olfactory system, odorants activate olfactory receptor neurons in the antenna. This signal is transmitted downstream to the antennal lobe (analogous to the vertebrate olfactory bulb) where it drives responses in cholinergic projection neurons (PNs) and GABAergic local neurons (LNs). The interaction between PNs and LNs transforms the sensory input received into complex patterns of activity distributed across ensemble of PNs that become the output of the antennal lobe circuit. Prior work has shown that information about the identity and intensity of an odorant is encoded by spatiotemporal PN activity patterns. While individual PN responses were perturbed by manipulating stimulus dynamics [11, 14], stimulus history [15, 16], and presence of background chemicals [7], the ensemble neural patterns can still allow recognition of odorants. Behavioral evidences also support this interpretation and reveal that odorants can be recognized independent of background cues [7] and stimulus history [15].

It is worth noting that prior studies examined neural response variability that arises due to each of these perturbations in isolation. In natural contexts, such interferences could occur independently or in conjunction with one another. Could robust odor recognition still be achieved? Particularly, can the variable neural responses be decoded in a manner that can simultaneously allow invariant odor recognition independent of all these perturbations? If so, what neural response features would be important for achieving this result? We sought to examine these issues in this study.

## RESULTS

### Stimulus dynamics, history and competing cues induce variations in PN responses

We began by examining how odor-evoked responses of individual projection neurons (PNs) in the locust antennal lobe were perturbed due to variations in how the odorant was encountered. Changes in pulse durations and inter-pulse intervals (stimulus dynamics), presence of other competing odorants (i.e. background vs. no background), and alterations in stimulus history (following termination of another cue) were all explored. A photoionization detector was used to characterize stimulus dynamics achieved using this delivery protocol (**Supplementary Figure 1**). For all odorants used, we found that the PID signals reached a steady-state level within 500 ms of pulse onset. Return to baseline was relatively slower and took ∼1 s or more in some cases. Pulses with less than a second of inter-pulse interval, or that were overlapping, had an additive effect on the photoionization detector. Overall, the stimulus delivery was robust, and a consistent steady-state level stimulus concentration was reached independent of how the odorant was delivered.

We recorded responses of eighty-nine PNs in the locust antennal lobe (n = 25 locusts). First, we examined the ability of individual PNs to robustly encode the identity of two ‘target’ odorants, hexanol (hex) and isoamyl acetate (iaa), that were encountered during the complex stimulation procedure (**Figure 1A-D**). In the entire ensemble of PNs that were recorded, we found that four PNs responded robustly to all encounters of the target odorants (**Figure 1A, B**; PN1 and PN2). Since these ‘reliable’ PNs were activated by both the target odorants (hex and iaa) they therefore did not provide discrimination between these two odorants. For all other PNs that were reliably activated during the first pulse of the target odorants, the responses during the subsequent encounters were either attenuated or completely suppressed due to the presence of other competing odorants and/or changes in stimulus histories (**Figure 1C, D**; PNs 3-8).

**Figure 1:**
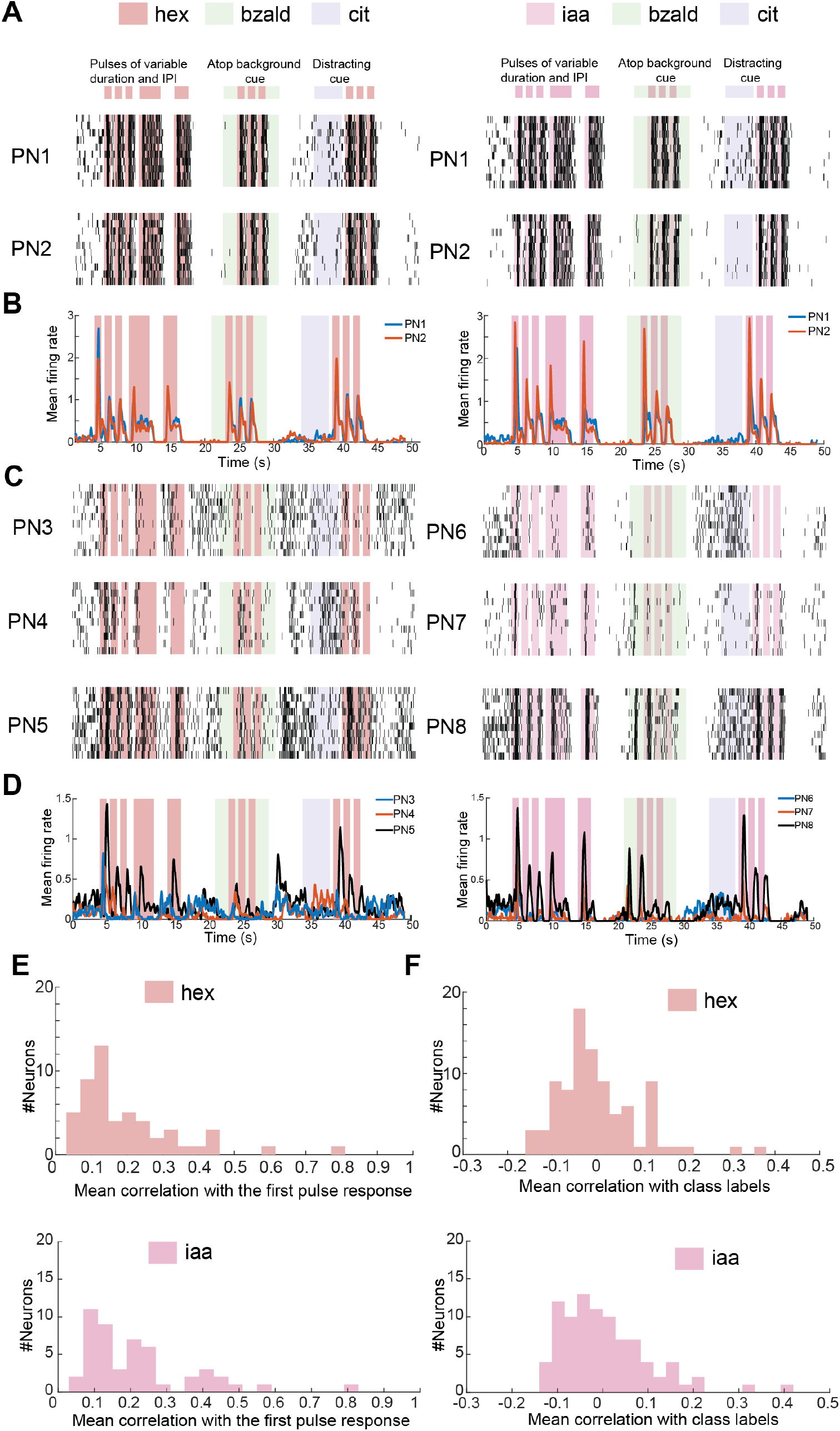
Individual projection neuron responses are highly variable. **(A)** Left plots, raster plots showing PN responses (PN1 and PN2) during a pulsatile presentation of a target stimulus (hexanol; *hex*) in back-to-back sequences of variable duration and inter-pulse intervals, atop a background cue (isoamyl acetate; *iaa*), following a distracting stimulus (citral; *cit*). Each black tick represents an action potential fired by the PN. PN responses are shown for 10 consecutive trials (10 rows). Right plots, similar plots as in the left, but the target stimulus was isoamyl acetate (*iaa*). Notice that these PNs responded reliably across all the presentations of both *hex* and *iaa*. **(B)** Firing rates of the two PNs (50 ms time bins; trial averaged) shown in **panel A** are now plotted as a function of time. While both the PNs responded strongly to the first pulse of the target odorant, the response diminished during later encounters of the same stimuli. **(C)** Similar plots as **panel A** but shown for three different PNs. Unlike the PNs shown in **panel A**, the responses evoked by the target odorant in these six PNs were highly variable. **(D)** Similar plots as in **panel B** but time histograms are plotted for the PNs shown in **panel C**. **(E)** Similarities between PN responses evoked during the first pulse of the target odorant with all other encounters were computed. For this quantification, PN response was first binned into 50 ms time bins and averaged across 10 trials. The first 1 s response following onset of each target odorant pulse was used to compare response similarity between different target odor encounters (i.e. 20-dimensional response vectors). For each PN, the mean similarity across odor pulses was determined, and the response similarity across PNs were then plotted as a distribution. Top and bottom plots reveal response similarity distribution for *hex* and *iaa*, respectively. **(F)** Similar plot showing correlation between the PN responses and the odor class label. The class label was set to ‘0’ corresponding to timebins when the target stimulus was not presented and set to ‘1’ for those timebins when the target stimulus was presented. Top and bottom plots reveal the correlation between individual PN responses and odor class labels for the two target odorants: *hex* and *iaa*.

To quantify the response variability observed at the level of individual PNs, we computed correlations between the PN response to the first pulse of target odorant and all the other introductions of the same chemical (see **Methods**). The distribution of these response correlations revealed that spiking activities during subsequent encounters of the target odorant had only a weak pattern match with the responses elicited during the very first encounter of an odorant (**Figure 1E**). Furthermore, when individual PN responses were correlated with an ‘odor label’ (i.e. a binary vector to indicate whether the target odorant was present or absent), the distribution was again centered around zero indicating that individual PN activities were not a reliable indicator of presence or absence of any target odorant.

### Variations due to changes in ambient conditions

Next, we examined whether changes in humidity conditions would further exacerbate the problem of robustly encoding odorant identity. For this purpose, we used the same stimulus delivery protocol but using either dry air (0% relative humidity (RH)) or humid air (100% RH) as the carrier stream. We again found that the four PNs that had robust responses to the target odorants also had reliable activity in varying humidity conditions (**Figure 2A, B**; PN1). However, for all other PNs in our dataset the responses were again variable in both dry and humid conditions (**Figure 2A, B**; PN9, PN 11). The overall distribution of response similarity between the first pulse and the subsequent encounters of the same odorant was low but comparable in both dry and humid ambient conditions (**Figure 2C**).

**Figure 2:**
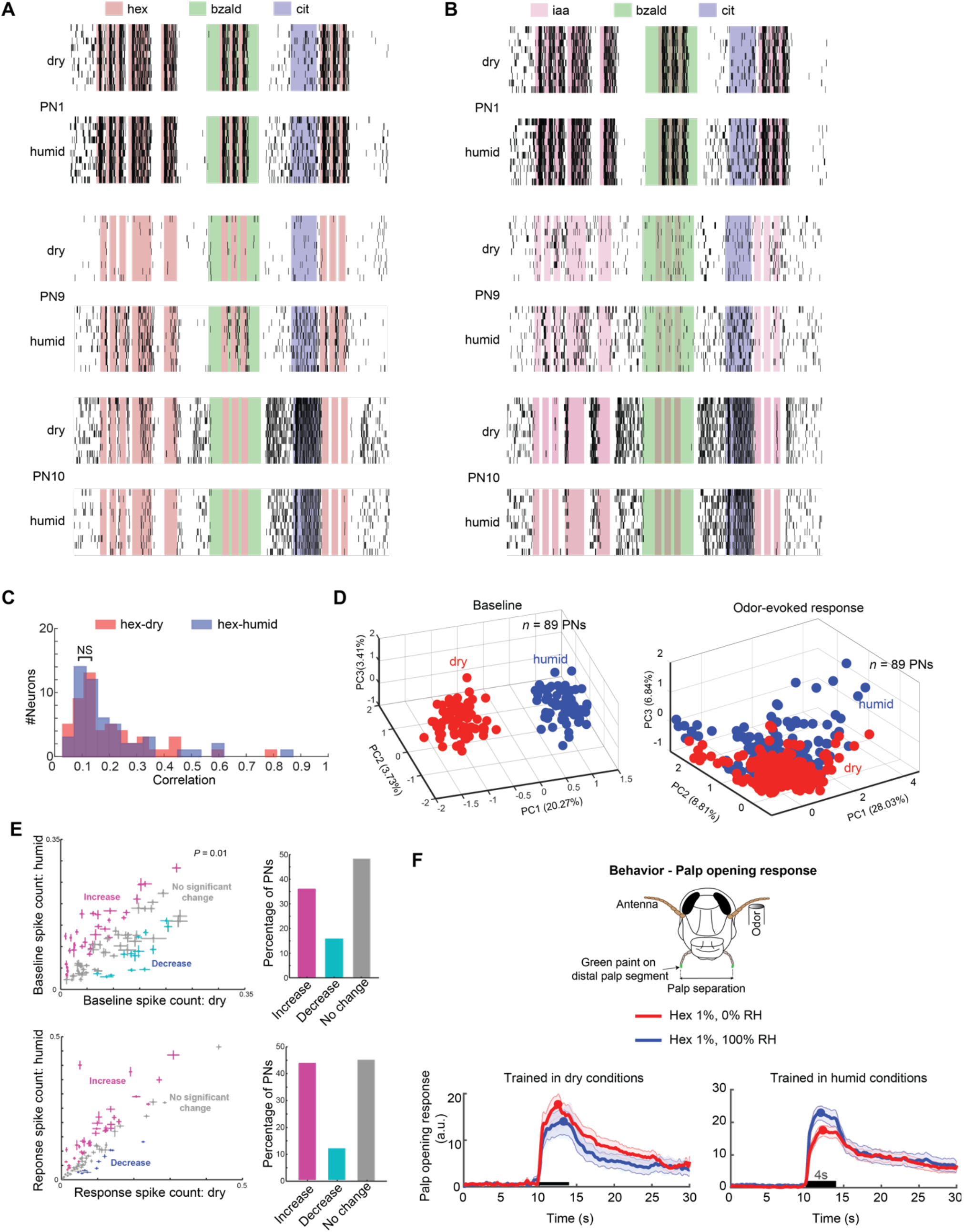
Spontaneous spiking activity and stimulus-evoked PN responses are both altered in different humidity conditions. **(A)** Similar raster plot showing PN responses to the stimulation protocol used in **Figure 1**. For each PN shown, the top and bottom plots reveal the spiking activity of the same PN between dry (carrier stream – 0% RH) and humid (carrier stream – 100% RH) conditions are shown. Note that changes in humidity levels of the carrier stream resulted in increase or decrease in spiking activity in individual PNs. **(B)** Similar plots as in **panel A** but showing PN responses to a different target stimulus (*iaa)*. Note that the same set of PNs are shown. **(C)** Similar plot as in **Figure 1D** but comparing response similarity between PN responses observed in dry and humid conditions. ‘NS’ indicates that the two distributions are not significantly different (Two-sample Kolmogorov-Smirnov test; *P* = 0.05). **(D)** Left, spiking activities of all 89 PNs during a 15 s pre-stimulus period (i.e. before odor stimulation begins) are shown after PCA dimensionality reduction. Spikes in individual PNs were binned in 200 ms non-overlapping windows and treated as vector components. Each colored sphere represents a 89-dimensional PN spike count vector along the first three principal components. Red and Blue colored spheres are used indicate the differences observed in the spontaneous, ensemble PN spiking activity in dry and humid conditions, respectively. Right panel shows a similar plot but comparing odor-evoked responses in dry and humid conditions (see **Methods**). **(E)** Top panel, Comparison of projection neuron spike counts during a 15-s pre-stimulus period (i.e. no odorant present). The *x* axis corresponds to spike counts in dry conditions. The *y* axis corresponds to spike counts in humid conditions. The mean ± s.e.m. over ten trials is shown for all PNs. Markers are colored to indicate significant increase (magenta), decrease (cyan), or no change in spike counts (paired t-test, *P* = 0.01, *n*=10 trials). On the right, bar plot is shown summarizing changes in baseline firings in individual PNs. Note that ∼50% PNs had changes in baseline activity. Bottom panel, similar plots as in top panels but comparing odor-evoked PN responses to hexanol in dry and humid conditions are shown and summarized. **(F)** Behavioral palp-opening response assay is schematically shown. Locusts were trained to associate the conditioned stimulus (CS; hex 1% v/v) with a food reward (US; grass). Subsequently, in an unrewarded testing phase, trained locusts opened their sensory appendages close to their mouths called palps to indicate successful recognition of the conditioned stimulus. Note that the palps were painted with a non-odorous paint and tracked to quantify behavioral palp-opening response. Locusts were trained in either dry or humid conditions but subsequently tested in both conditions (see Methods for more details). Median palp opening response ± s.e.m. (*n*=20 locusts) is shown for the testing phase trials. Note that POR responses to the conditioned stimulus in shown for both dry and humid conditions.

Between dry and humid conditions, we found that both spontaneous activity and odor-evoked responses varied in ∼ 50% of PNs recorded (**Figure 2D, E**). While most PNs had increased activity in the humid conditions, a smaller fraction had higher baseline activities and odor-evoked responses in dry conditions. These results suggest that changes in ambient conditions (i.e. dry or humid) can alter both spontaneous activity (i.e. initial state of a dynamical system), and odor-evoked neural response dynamics at the individual and at the ensemble level in the antennal lobe.

These physiological results raise the following question: can locusts recognize an odorant independent of the changes in humidity conditions? To examine this, we trained locusts in an appetitive conditioning assay. In this paradigm, each locust was trained by presenting an odorant (conditioned stimulus; hexanol at 1 % v/v concentration) in dry conditions followed by a grass reward (unconditioned stimulus) in each training phase trial. Following six training trials, we tested the ability of the trained locusts to recognize the conditioned odor stimulus in unrewarded test phase trials. The opening of palps (sensory appendages close to the mouth; palp-opening response or POR) in anticipation of grass reward, following the introduction of the conditioned odor stimulus was interpreted as successful recognition of the odorant. After the training phase in dry condition, we examined the ability of locusts to recognize the conditioned stimulus presented either in dry or humid conditions. Our results show that locusts opened their palps to all the introductions of the conditioned stimulus in both dry and humid conditions (**Figure 2F**). The performance was near identical indicating robust odor recognition that was invariant with respect to changes in ambient conditions. Similar results were also obtained when locusts were trained in humid conditions and tested in both dry and humid conditions (**Figure 2F**). These results indicate that locusts can recognize trained odorants invariant to changes in ambient conditions.

### Bayesian optimal Boolean decoder for robust odor recognition

Our results indicate that odor-evoked neural responses vary with stimulus dynamics, history, the presence of competing cues and changes in ambient conditions. However, the behavioral recognition (i.e. the POR) remained invariant to such perturbations (**Figure 2**; also refer prior results on background [7] and history invariance [15]). Given this discrepancy between variability in neural encoding and robustness in behavioral output, we sought to determine whether a neural decoder could be designed for robust odor recognition.

To investigate this issue, we regarded the ensemble activity across the 89 PNs recorded in a 50 ms time bin as a snapshot of high-dimensional neural activity (i.e. 89-dimensional firing rate vector). To design the simplest decoder, we assumed that a linear Bayesian logistic regression scheme with binary components (i.e. {‘0’ or ‘1’} weights) would be sufficient to allow robust recognition (**Figure 3A**; see **Methods**). Note that spontaneous and odor-evoked responses observed in a separate set of trials that included solitary exposures to two target odorants were used to determine the binary weight vector (**Supplementary Figure 2**). In essence, multiplying the PN high dimensional activity vector with the binary vector (dot product) would result in combining a spiking activity of all individual PNs that were assigned a weight of ‘1’ and ignoring responses in all other PNs.

**Figure 3:**
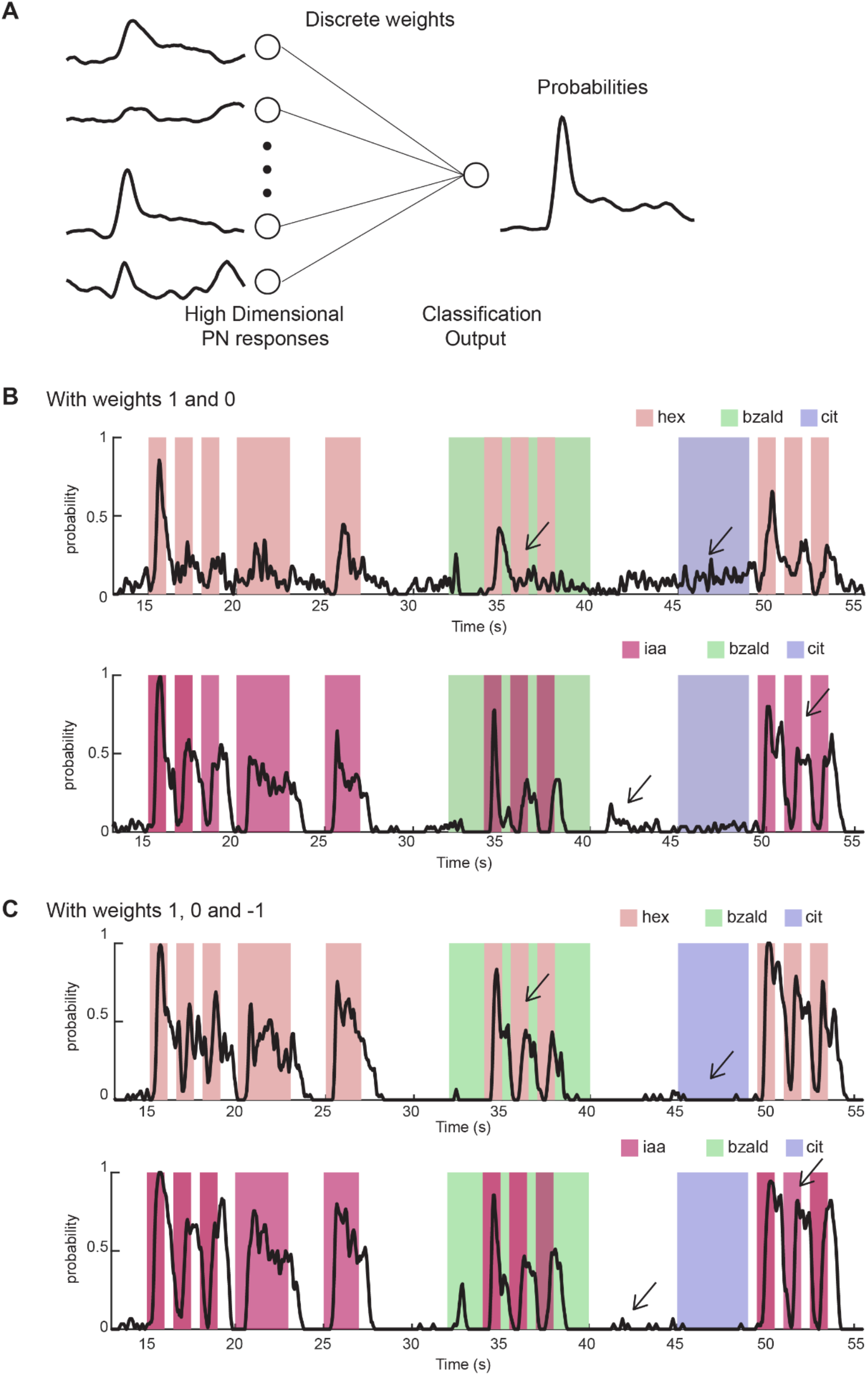
Robust odor recognition with a Bayesian logistic regression approach. **(A)** The decoding approach is schematically shown. Response of each PN was binned in 50 ms timebins. The spike counts across all 89 PNs in a given timebin became input to the Bayesian logistic regression classifier. This classification scheme had 89 free parameters corresponding to the weight assigned to each PN. The approach was constrained such that the weights could only assume either binary {‘0’ or ‘1’} or ternary values (‘-1’ or ‘0’ or ‘1’). The outputs from the classifier the target odorant classification probabilities (see **Methods** for more details). **(B)** The output of logistic regression scheme with binary weights are shown. Note that for each time bin, the classifier output indicates the probability that the target stimulus was present (i.e. averaged across trials; refer Supplementary Figure 3 for trial-by-trial results). Top panel shows the classification probabilities for trials where *hex* was the target stimulus, and the bottom panel shows the classification probabilities when using *iaa* as the target stimulus. **(C)** Similar to **panel B**, but for a logistic regression classifier with ternary weights {‘-1’ or ‘0’ or ‘1’}. Note that both increased recognition of target stimulus and reduction of false positives were both observed (arrows).

Odor recognition (trials-averaged) results from this binary classification scheme for the complex, pulsatile stimulus delivery are shown in **Figure 3B (**refer **Supplementary Figure 3** for trial-by-trial classification results). As can be noted, both target odorants were detected and recognized during all the introductions. In general, recognition performance was better for *iaa* than *hex* as more PNs in our dataset were activated by *iaa*. The best recognition was observed in the very first odor pulse and the performance progressively dipped during subsequent encounters. Recognition of the target odorants was harder atop a background, particularly during the second and third pulse of *hex* atop *bzald (***Figure 3B**; *arrow head)*. Overall, these results indicated that a very simple Boolean classifier can achieve better than chance recognition of the target odorants.

Next, we relaxed the binary constraint on the weight vector and allowed each component to assume a value of ‘1’, or ‘0’, or ‘-1’. This minor modification now allowed penalizing spiking activities in a subset of PNs. Results from this ternary classification scheme are shown in **Figure 3C**. This modification improved recognition performance significantly. Both target odorants could now be recognized robustly, and recognition in the presence of backgrounds was also improved.

To understand the logic behind assignment of weights to each PN using the ternary approach, we first classified the responses of each PN to the target odorant as: ON response, OFF response, or no response (**Figure 4**). ON responsive PNs were activated during target odor exposure, whereas OFF responsive PNs were usually inhibited during odor exposure but became active after termination of the stimulus (see Methods). Not surprisingly, we found most ON neurons were assigned a ‘1’ weight and most non-responsive ones were assigned a ‘0’ weight. Notably, most OFF responsive PNs were assigned a ‘-1’ weight. This result was observed in weight vectors used for classifying both target odorants.

**Figure 4:**
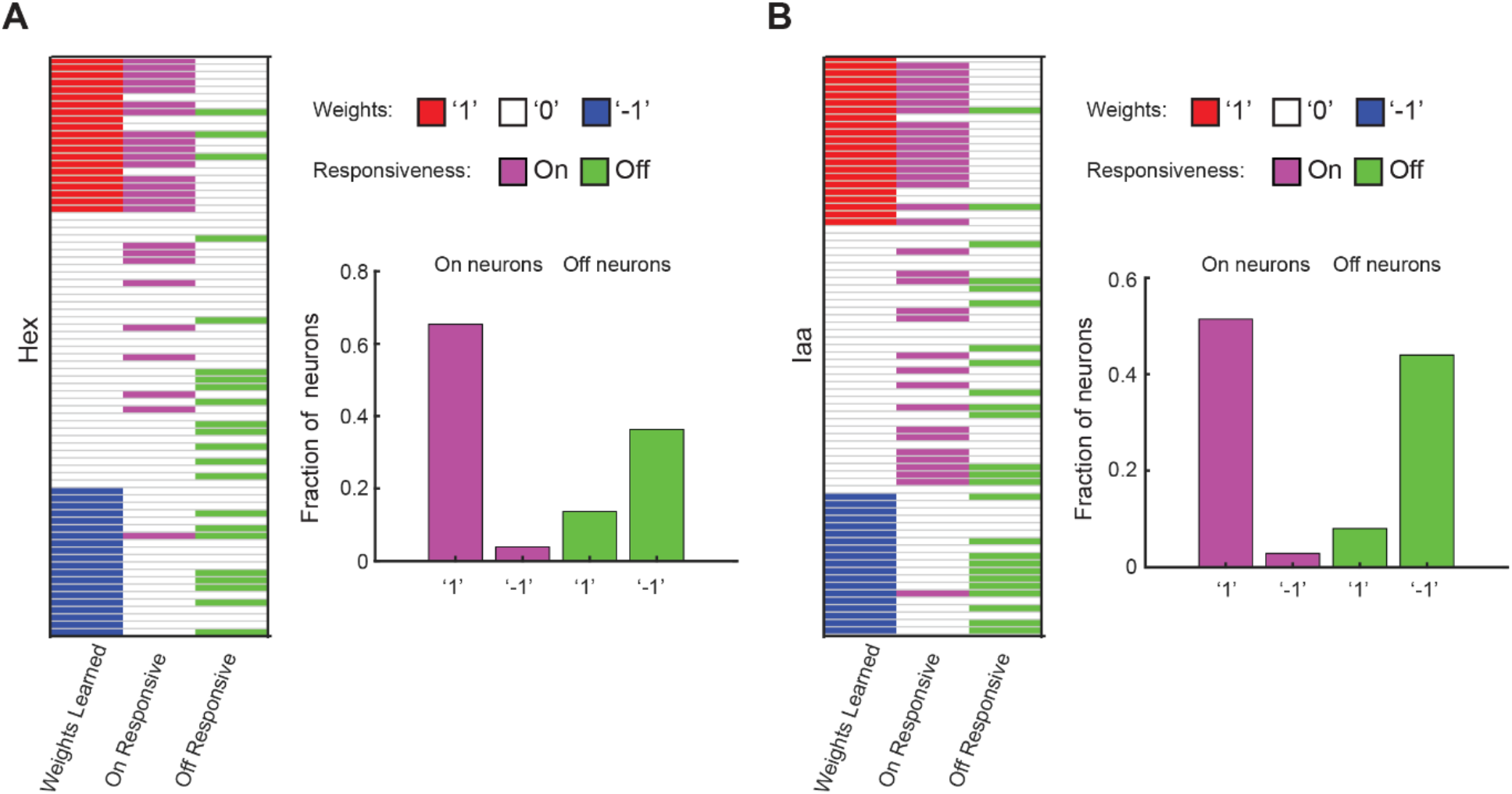
Logisitic regression weights assignment for ON and OFF neurons. **(A, B)** Weights that were assigned to each PN in logistic regression classification approach with ternary constrains are schematically shown. The first column shows the ternary weights learned using Bayesian framework as a heatmap. Neurons assigned ‘1’ weights are labelled in red. Neurons assigned ‘-1’ weights are labelled in blue. Neurons assigned ‘0’ weights neurons are labelled white. The second column reveals whether each PN was activated during the presentation of the target stimulus i.e. ON responsive (*hex* for panel A and *iaa* for panel B). The third column reveals whether each PN was activated after the termination of the target stimulus i.e. OFF responsive. The percentage of overlapping are summarized in bar plots. Panel **A** shows Hex as target stimulus and panel **B** shows Iaa as target stimulus. Bar plots compare the following four categories: fraction of ON neurons assigned +1 weight, fraction of ON neurons assigned -1 weight, fraction of OFF neurons assigned +1 weight and fraction of OFF neurons assigned -1 weight.

Taken together, our results indicate that although individual PN responses vary unpredictably, a simple linear neural decoding scheme is sufficient to achieve invariant odor recognition.

### Robust recognition with a ‘ON minus OFF’ classifier

Our results indicate that Bayes optimal logistic regression approach weighted most of the ON responsive neurons with a ‘1’ weight and OFF responsive neurons received a ‘-1’ weight. We wondered whether this relatively simple approach of directly summing up firing rate in ON neurons and subtracting the summed contribution from the OFF neurons would make a successful classifier. We again classified each PN as a ON responsive or OFF responsive based on its response during the training phase trials that only included solitary exposures of the target odorants (**Supplementary Figure 2**). The classification approach had two components: an ON classifier that summed the contribution of all ON responders (achieved using a binary weight assignment of ‘1’ to all ON neurons and ‘0’ to all others) that was followed by a thresholding step, and an OFF classifier that summed the contribution of all OFF responders (achieved using a binary weight assignment of ‘1’ to all OFF neurons and ‘0’ to all others) followed by another thresholding step. The outputs of these two classifiers were combined by computing the difference between the ON classifier output and the OFF classifier output. Only when the resulting output was positive, the target stimulus was predicted to be present (i.e. a rectifying linear unit or a ‘ReLU’). Note that the only two free parameters in this approach are the classification thresholds that were set to minimize false positives during pre-stimulus, while maximizing true-positives in the training data. As can be observed robust recognition of the target odorants can be achieved using this extremely simple approach (**Figure 5 A-C**). Our results indicate that while the ON neurons alone could help recognize the target odorants, they would also trigger a few false positives (**Figure 5A**). Inclusion of OFF neurons helped eliminate some of these false positives and improved recognition performance. In sum, our results indicate that a simple but generic approach can achieve invariant odor recognition in this model system.

**Figure 5:**
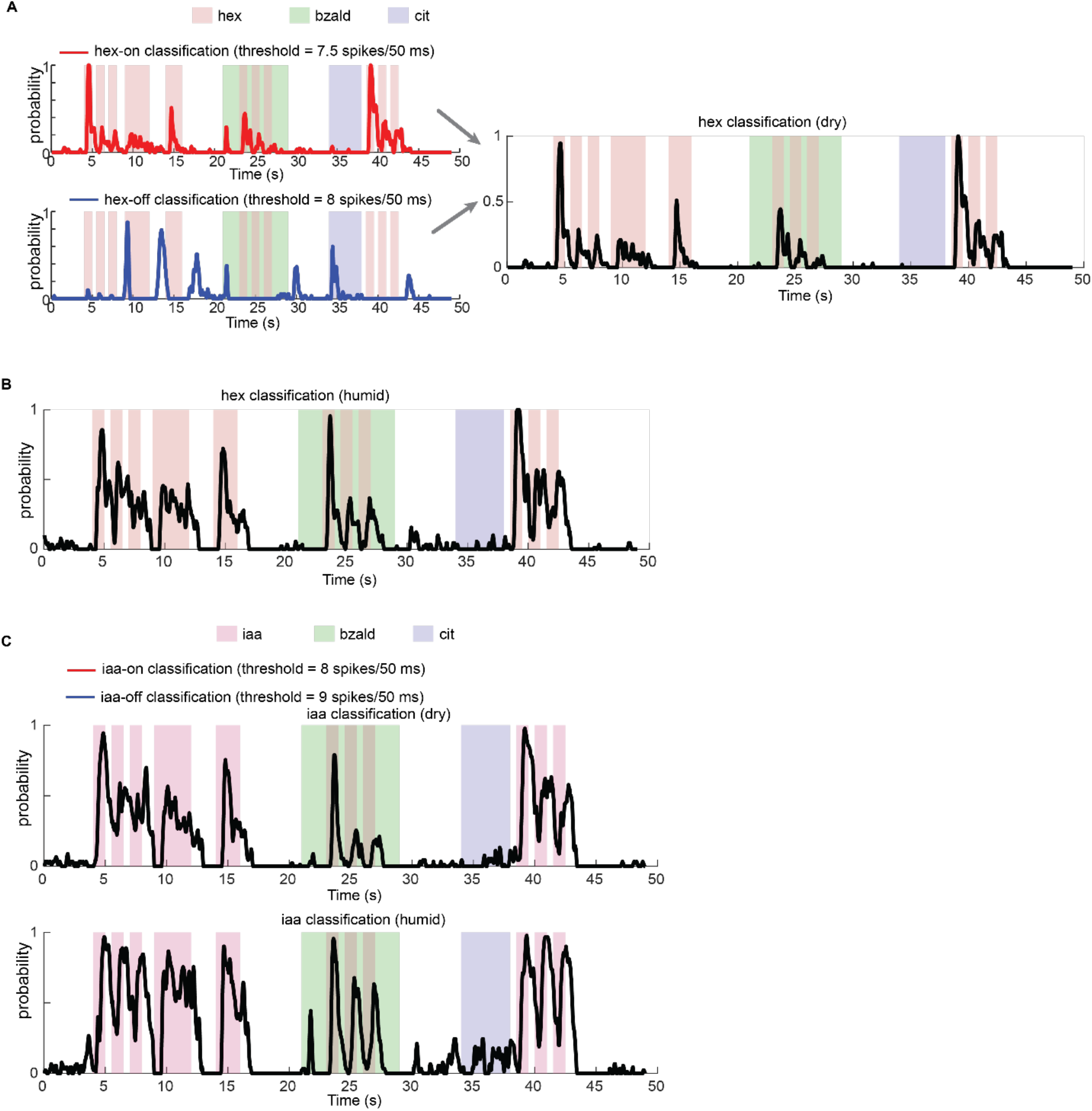
Decoding stimulus identity with a ‘ON minus OFF’ classifier. **(A)** Top, classification probability for *hex* as a function of time using a ON classifier is shown. ON responsive neurons were first found using solitary presentation of *hex* (see Methods; **Supplementary Figure 2**). In other words, a weight of ‘1’ was assigned to the ON responsive PNs and 0 to the rest. At each time bin, PN firing rates from ON responsive neurons were summed and thresholded to classify each timebin. Trial-by-trial classification results were averaged to generate classification probabilities and plot them as a function of time. Bottom panel, similar plot but showing classification results using OFF classifier that predicts the absence of the target stimulus using OFF-responsive neurons. Thresholds used for both these classifiers are shown at the top of each panel. Right, ON and OFF classifications were merged with a rectified linear unit (ReLU) as an activation function that outputs non-zero only if the ON classifier output is greater than the output of the OFF classifier and zero elsewhere. **(B)** Classification probabilities predicted using a ‘ON minus OFF’ classifier is shown for hexanol stimulation in humid condition. **(C)** Similar plots as in **panels A and B** but showing classification probabilities for *iaa* predicted using ON minus OFF classifier. Top and bottom sub-panels show robust recognition results in dry and humid conditions, respectively.

### A generic binary neural network inspired by the insect olfactory system

We wondered whether the simple approach for achieving invariance in the locust olfactory system can be extended to create a general-purpose pattern recognition neural network. Note that the goal is to develop an artificial neural network (ANN) that has a similar architecture to the insect olfactory system and uses binary values ({-1 and 1} or {0 and 1}) as learned parameters/weights for connections between individual nodes (**Figure 6A**). Most ANNs use real-valued parameters that are unconstrained and routinely learned through back-propagation algorithm. However, with a Boolean or ternary weights constraint, backpropagation with regular gradient descent would not be feasible (**Supplementary Figure 4; see Methods**). To overcome this issue, we reformulated this problem as one of fitting a Bayesian model and choosing a Boolean distribution as a prior distribution for ANN weights. As direct inference of weights using the posterior distribution would be intractable, we used a variational inference approach to approximate the posterior with a lower bound. An unbiased estimator of the derivative of lower bound can be found by relaxing and reparametrizing the weights using Gumbel-Softmax variables (**Supplementary Figure 4; see Methods**).

**Figure 6:**
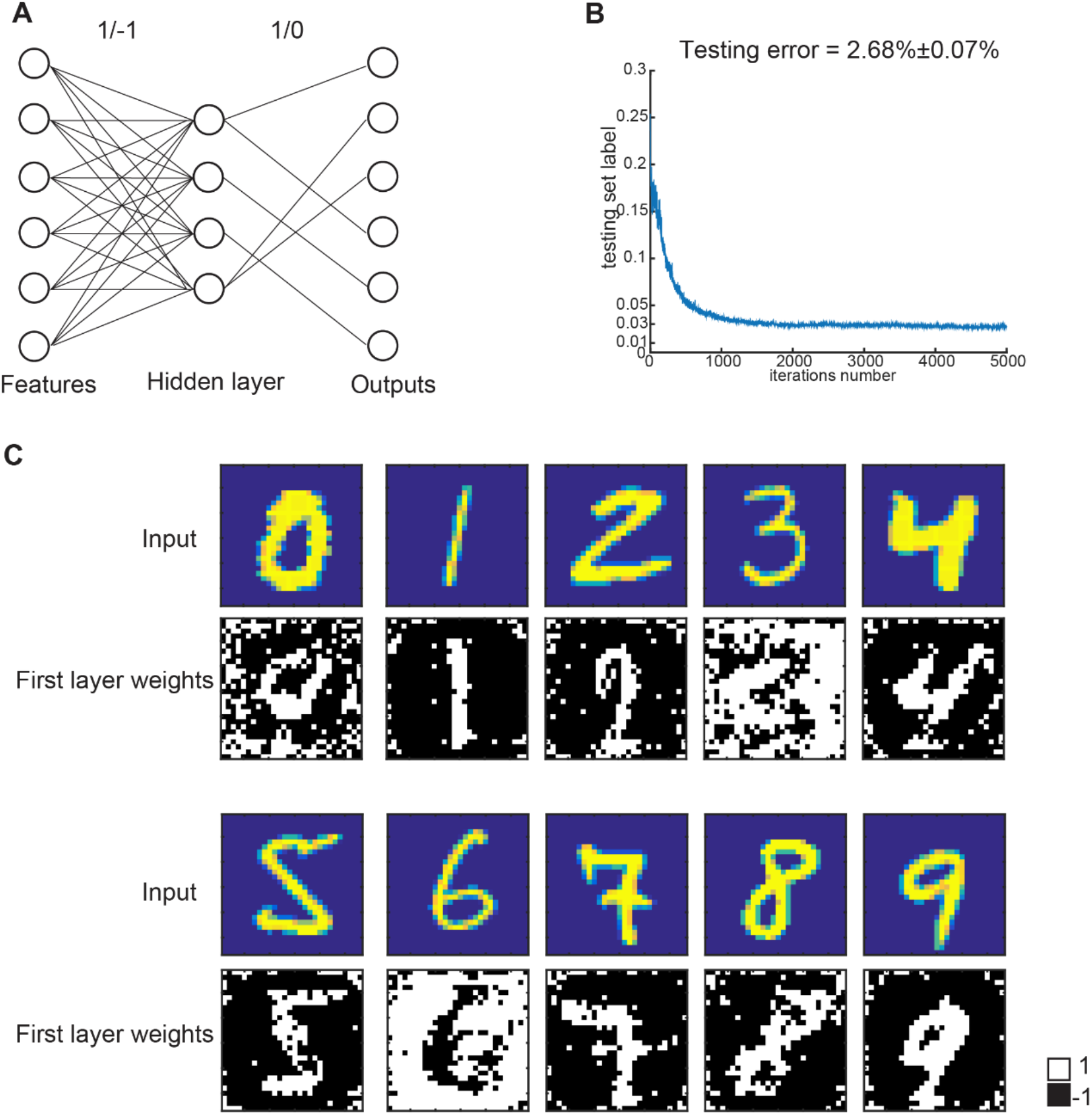
Boolean neural network for non-olfactory pattern recognition. **(A)** Schematic of a feed-forward neural network model. Note that all the weights were constrained to be discrete. The weights from the first layer to hidden layer were constrained to be either 1 or -1. The weights from hidden layer to output layer are constrained to 1 or 0. **(B)** The learning curve of network shown in panel **A**. **(C)** Representative learned weights connecting the input layer with the hidden layer neurons are shown. To compare with the input patterns, the weights were reshaped to match the size of input image. In each sub-panel, the top image shows a ‘close’ input digit and the bottom image shows the representative weights. Note that weight assignment mimicked ON pixels to+1 (digits 0, 1, 2, 4, 5, 7, 8, 9) and OFF pixels to -1 or *vice versa* (digits 3, 6).

An advantage of this Bayesian neural network architecture is that it can be simply scaled to multiple layers in a neural network. Additionally, we can train the neural network with different layers that take different combinations of weights (for example, binary or ternary), but are still limited to a set of discrete weights. This is particularly advantageous when a desired outcome can be obtained from an interpretable neural network model.

To examine how well a neural network with only binary weights can perform, we trained a fully-connected neural network with one hidden layer on the MNIST dataset (**Figure 6**). We constrained the first layer with weights {1, -1} and the second layer with weights {1, 0}. After training to classify the ten digits, we found that the loss function had converged within a similar window of iterations compared to a regular neural network with real-valued weights (**Figure 6B**). Next, we wondered how this simple neural network compares with state-of-the-art deep neural networks when it comes to image classification. The performance of the trained binary weights neural network on a held-out dataset revealed an error rate of 2.68%, only modestly poorer than the benchmark performance of 1.8% for an ANN with a similar architecture but with real valued weights [17]. This result indicates that constraining the weights in a layer of neural network with just two values only results in a minimal drop in the performance.

As mentioned earlier, the binary valued ANN allowed us to understand what features were extracted and how they were used to improve classification performance. We found that the hidden-layer neurons performed two types of feature extraction (**Figure 6C**): a subset that weighted ‘ON pixels’ for a digit with a weight of ‘1’ and ‘OFF pixels’ with a weight of ‘-1,’ and another subset that weighted the pixels in the opposite fashion (i.e. ‘-1’ for ON pixels and ‘1’ for OFF pixels). It is worth noting that this approach is strikingly similar to the one we used for robust recognition of the odorant in this study. In sum, these results indicate that a binary neural network, inspired based on results from the insect olfactory system, performed relatively well as a general-purpose pattern recognition algorithm.

## DISCUSSION

We examined how invariant recognition of odorants can be achieved in a relatively simple locust olfactory system. Our results indicate that individual PN responses can vary with one or several of perturbations we studied, including, stimulus dynamics, repetition, stimulus history, presence of background odorants and changes in humidity conditions. Nevertheless, at the population level information regarding the odorant identity was still robustly encoded, and a simple classification scheme was sufficient to extract the relevant information. The classifier essentially boiled down to adding the contribution of PNs that were strongly activated when the odorant was presented (ON neurons) and subtracting the contribution of PNs that were activated after the termination of the odorant (OFF neurons). We generalized this simple scheme to develop a Boolean neural network that was highly interpretable and performed well in non-olfactory pattern recognition tasks. In sum, these results indicate a neural basis for achieving sensory invariance.

We found that not all neurons were perturbed and only a small subset (4/89 PNs) of them responded reliably to all introductions of the target odorants. While these neurons allowed robust detection of the target odorants, they were not specific and responded to both the target odorants examined in this study. Therefore, it is possible that an approach based on a single or on a small subset of neurons encoding for a stimulus under all conditions may not be fault-tolerant.

Prior publications [7, 11, 14-16] had also found individual neurons to be unreliable, but found that robustness emerged at an ensemble level. However, our prior results indicated that odorants delivered atop different background cues generated ensemble responses that only partially overlap across conditions [7]. Furthermore, features of ensemble responses also varied unreliably. In other words, even at the ensemble level, there was not a single feature that remained consistent when the odor-evoked responses were minimally perturbed. How then could sensory invariance be achieved?

Given the complexity of individual PN response changes, we did not expect a linear classifier, with binary or ternary weights constraints, could provide robust recognition. The goal for constraining the decoding scheme was two-fold: interpretability of the approach taken and determining the simplest possible approach. Our results indicate that not only did such a scheme exist, but it exploited a simple stratagem. We found that the weights assigned by the Bayes optimal logistic regressor assigned ‘+1’ weights to ON neurons, ‘0’ weight to non-responders, and ‘-1’ weight to OFF neurons (i.e. a ‘ON minus OFF’ classifier).

If ensemble responses features varied across conditions how did this ‘ON minus OFF’ classification approach achieve invariance? It is worth breaking this classification scheme into its two components: the ON component and the OFF component. Assigning ‘+1’ to most strongly responding ON neurons and setting a recognition threshold that is less that this sum allows the classification scheme to be flexible. Interpreted differently, this indicates that an odor can be recognized as long as a subset of strongly responding ON neurons are activated so that their sum reaches the threshold. The composition of this subset can change across conditions thereby allowing this approach to be more flexible.

What then is the contribution of the OFF component of the classifier? In an earlier study[8], we found that OFF responses were better at predicting when the behavioral response to a conditioned odorant terminated. Here, in this study we found that the OFF component increased separability between activation patterns of different odorants. This enhanced discrimination between odorants and thereby reduced false positives.

Our results indicate that not only the odor-evoked responses but also the spontaneous activity can change across conditions. Earlier studies have argued that the antennal lobe neural network can be viewed as a non-linear dynamical system [6, 18, 19]. Under this perspective, our results indicate that both the initial conditions and odor-evoked response dynamics can vary across conditions. Yet at direct odds with our neural data, we find the behavioral recognition is robust even during these drastic changes. No detectable differences in response latency, intensity or duration were found. Hence, our results indicate that the rules for translating neural responses and their dynamics to generate appropriate behavioral output needs further investigation.

Given the recognition performance in the olfaction dataset, we finally explored how well this scheme would generalize to detecting and discriminating patterns in other domains. To examine this, we developed a Boolean neural network directly inspired by the architecture and results from the locust olfactory system and applied it to classifying patterns in the well-studied MNIST digits database[20]. Although our network was highly constrained, our results were notably comparable to other state-of-art techniques [17]. In addition, the Boolean weights made the features extracted by each hidden unit highly interpretable. Taken together, these results indicate that the neural circuits in the insect olfactory system delicately balance discrimination between odorants with flexibility necessary for robustness and fault tolerance to achieve sensory invariance.

## Acknowledgements

We thank members of the Raman Lab (Washington University in St. Louis) for feedback on the manuscript. We thank Pearl Olsen for insect care. This research was supported by NSF CAREER Award (Grant #1453022) and NSF CRCNS (Grant#1724218) to B.R.

## Author contributions

SN, LZ and BR conceived the study and designed the experiments/analyses. SN performed all the electrophysiology experiments and collected the data. RC collected the behavioral data. LZ developed the Bayesian logistic regression approach and the Boolean Neural Network. SN, LZ and BR wrote the paper taking inputs from all the authors. BR supervised all aspects of the work.

## METHODS

### Odor stimulation

Neat odor solutions (Sigma-Aldrich) were diluted in mineral oil to their 1% concentration by volume (v/v). The diluted odor solution was placed in a 60 ml sealed glass bottle with separate inlet and outlet lines. A pneumatic pico-pump (WPI Inc., PV-820) was used to displace a measured volume of the odor-bottle headspace (0.1 L min^-1^) that was then injected into a desiccated and filtered carrier air stream (0.75 L min^-1^) directed towards the locust antenna. A vacuum funnel was placed right behind the locust antenna to provide a constant flux and ensure removal of the delivered odor vapors. Each odorant/combination-of-odorants was presented in a pseudorandom manner (blocks of 10 trials). The inter-trial interval was 60 s long for solitary odorant exposures, or 80 s for the lengthy protocol shown in **Figure. 1** and **Supplementary Fig. 1**. The following odorants were used in this study: hexanol (*hex*), isoamyl acetate (*iaa*), benzaldehyde (*bzald*), and citral (*cit*).

### Electrophysiology

Young locusts (*Schistocerca americana*) with fully developed wings (post fifth instar) of either sex were selected from a crowded colony. Locusts were immobilized with both antennae intact, and then the brain was exposed, desheathed, and continually perfused with locust saline as demonstrated previously [11, 21, 22]. *In vivo* extracellular recordings from the antennal lobe were performed using 16-channel, 4×4 silicon probes (NeuroNexus). Electrodes were gold plated such that their impedances were in the 200 – 300 kΩ range. The extracellular signals were acquired using a LabView data acquisition system. Raw extracellular signals were collected at 15 kHz sampling rate, amplified at 10 k gain using a custom made 16-channel amplifier (Biology Electronics Shop; Caltech, Pasadena, CA), filtered between 0.3 to 6 kHz prior to spike sorting.

### PN spike sorting

Spike sorting was done using a conservative approach described in earlier works [7, 9, 23]. In brief, the following criteria were used for the single-unit identification: cluster separation > 5 noise s.d., number of spikes within 20 ms < 6.5%, and spike waveform variance < 6.5 noise s.d. Using this approach, a total of 89 PNs were identified from 25 locusts.

### PID experiment

We used a fast photo-ionization detector (miniPID, Aurora Scientific) to characterize the dynamics of stimulus delivered. Raw data were amplified (gain = 5) and acquired at 15 kHz sampling rate using a custom LabView data acquisition program (**Supplementary Figure 1A**).

### Behavior experiments

Locust behavior experiments were performed following previously published methods [8, 9, 24]. In brief, adult locusts of either sex were starved for 24 hours and then immobilized in plastic syringes with an opening cut to ensure the locust antennae, head, and mouthparts were visible and accessible. The eyes were closed with a black tape and the distal segments of the maxillary palps (mouthparts) were painted green with a zero-volatile organic compound paint (Valspar Ultra) for image tracking purposes. The training sessions began approximately an hour after the palps were painted to allow proper drying and to allow the locusts to acclimatize to being immobilized and have paint on their palps.

Training was performed in either a 0% or 100% relative humidity (RH) carrier air stream at 750 sccm. To achieve 100% RH, the dry air stream (i.e. 0% RH) was diverted through a water bubbler to be completely humidified. 1% Hexanol was used as the conditioned stimulus (CS) in training and organic wheat grass was the unconditioned stimulus (US). Odor pulses were presented at 100 sccm in addition to the air. To clear delivered odorants, a vacuum line was placed behind the locusts. A video camera (Microsoft Webcam) was used to record the locusts’ palp opening response (POR) at 30 frames per second. Odor delivery and video data acquisition were automated using custom written LabVIEW 2009 (National Instruments, Austin, TX) programs for precise testing conditions.

Locusts were trained over 6 training trials, spaced 10 minutes apart, in which the CS was presented for 10 seconds and the US was presented 5 seconds after the start of the CS for approximately 10 seconds. Only locusts which ate wheat grass in at least four out of the six training trials and performed a satisfactory palp-opening response (POR) in at least three of the six trials were retained for the testing phase (67% of the locusts in 100% RH (20/30) training and 73% of the locusts in 0% RH (22/30) training were retained using this criterion). Also, note that 2 locusts trained in the dry condition were later removed due to poor palp-tracking.

In the testing phase (results reported in **Figure 2**), locust PORs were collected for a total of 6 trials. The 6 trials were a pseudo randomized combination of 0.1%, 1%, and 10% hexanol being presented for 4 seconds in the presence of 0% or 100% RH air. The inter-trial delay was set to 25 minutes.

Locusts were kept on a 12 h day – 12 h night cycle (7 am – 7 pm day). All behavioral experiments were performed between 9 am – 3 pm.

### Principal component analysis

We used a linear principal component analysis to visualize high-dimensional PN spike counts. The spike counts observed for each PN in 50 ms non-overlapping time bins were binned to generate a n-dimensional vector of neural activity for each timebin (n = 89 PNs). The high-dimensional PN spike count vector was projected onto the top three eigenvectors of the data covariance matrix for purposes of visualization. These low-dimensional representations of PN activity were color coded depending on whether the target odorant was presented during that timebin (**Figure 2D**).

### PN response characterization

For ‘ON minus OFF classifier’, we classified projection neurons as ON-responsive if the spike counts in any time bin during the stimulus presentation exceeded mean + 6.5 s.d. of pre-stimulus activity (2 s window just before onset of any stimulus). Similarly, a PN was regarded OFF-responsive if it met the same criterion in a 4 s window after the termination of the stimulus (0.5 s to 4.5 s after stimulus termination. Note that a 500 ms window immediately after the termination of the odorant pulse was ignored as it confounded both ON and OFF responses. All PNs that did not meet either of these criteria set for ON or OFF responders were regarded as ‘non-responders’.

### Discrete weights neural network

Let us consider a classification problem with the samples {***x***_*n*_, *y*_*n*_}, where ***x***_*n*_ denote input neural activity vectors in *R*^*n*^, and *y*_*n*_ represents class labels (target odor present or absent). Our definition of discrete weights neural network builds on a Bayesian formulation. The joint distribution of the model can be written as:

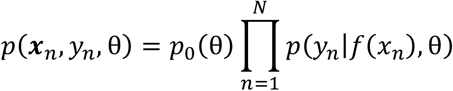

where θ represents weights, *p*_+_ θ is the prior distribution of weights, *f x*_*n*_ represents the final output of the neural network. Binary weights constraint can be achieved by choosing θ ∈ {-1,1}(The framework is flexible such that it can be used to train the with any combinations of discrete weights, with (−1, 0, 1) or (1, 0)). Using this probabilistic framework, we can now find the posterior distribution of the network weights *p* θ *x*_*n*_, *y*_*n*_ and use its statistics for future predictions. Since we constrained our weights to be discrete, a direct inference of weights’ posterior distribution would be intractable. We approximated the posterior distribution using variational inference (an approximation distribution *q*_*φ*_ θ parameterized by the variational parameters *φ*). We obtained the variational parameters by minimizing the KL divergence between the posterior distribution *p* θ *x*_*n*_, *y*_*n*_ and the approximate distribution *q*_*φ*_ θ, which is equivalent to maximize the evidence lower bound *L q*.

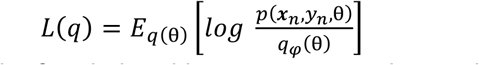

We followed the framework of variational bayes autoencoder to obtain an unbiased Monte Carlo gradient estimator of lower bound with respect to variational parameters. Furthermore, to address the difficulty of discrete weights, we sought to relax and reparameterize the weights θ using Gumbel-Softmax variables as proposed by [25-27] :

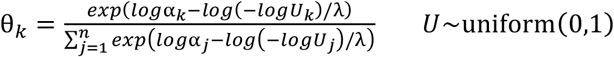

Then the gradient can be approximated unbiasedly by: 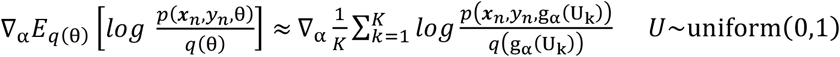

During the training phase, we can utilize the gradient information obtained by regular back-propagation algorithm to update the discrete weights.

For recognizing digits in the MNIST dataset (**Figure 6**), we trained a two-layer network with the constraint that the network weights were discrete. We followed standard stochastic gradient descent to update the parameters of the approximate distribution (see above) and adaptively choose the step size using ADAM optimizer.

The classification results shown in **Figure 3** was also obtained from discrete weights networks using the same training procedure. However, here only one layer of feedforward layer was used (see **Figure 3A**). To construct the training dataset, the neural responses to the solitary presentation of *hex* (or *iaa*) were used. The network output corresponded to the predicted class label: target odorant absent or target odorant present. Two networks were trained for recognizing *hex* or *iaa* (**Figure 3)**.

### ON minus OFF classifier

PN responses to 4 s long solitary presentations of target stimulus (hex or iaa; **Supplementary Figure 2**) were used to determine ON responders and OFF responders (using the same criteria discussed in PN response characterization above). For the ON classifier, a binary weight vector was obtained by assigning all ON responsive PNs a weight on ‘1’ and assigning all other PNs a weight of ‘0.’ PN responses were then linearly combined at each time bin by computing a dot product with the binary vector constructed. The summed activity was then compared to a threshold to classify ‘target present’ or ‘target absent’. Note that the threshold is a free parameter and needs to be determined to minimize false-positives during pre-stimulus period, while maximizing the true-positives during the stimulus. During testing, all the ten trials were used to quantify performance and the trial-averaged classification probabilities were computed as a function of time (shown in **Figure 5A**).

For the OFF classifier, the same strategy was used to obtain trial-averaged classification using a binary weight vector that assigned only the OFF responsive neurons a weight of ‘1’ (**Figure 5A**).

Finally, the difference of the two classifier’s output, i.e ON classifier output – OFF classifier output was computed for each timebin and passed through the rectifier (i.e. max(0, x)) and shown as the final result of ‘ON-minus-OFF’ classification approach.

## Supplementary Figure Captions

**Supplementary Figure 1:**
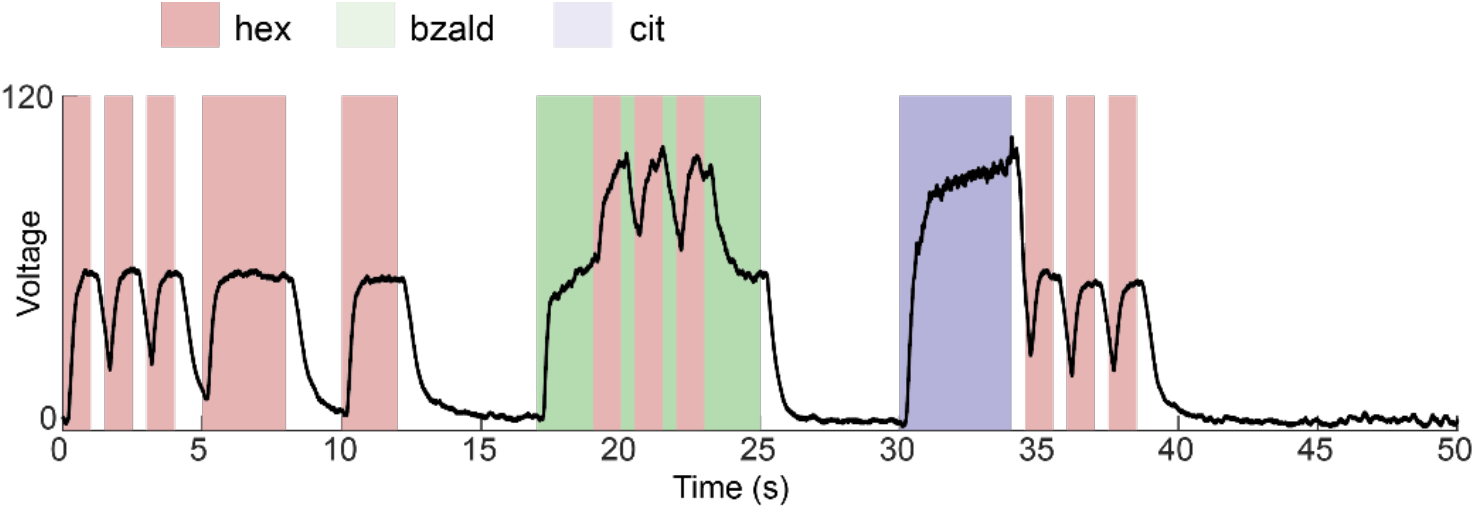
Voltage output of a miniPID (Aurora Scientific) is averaged (n = 3 trials) and plotted as a function time to characterize the stimulus delivery protocol. Same color convention as in **Figure 1**.

**Supplementary Figure 2:**
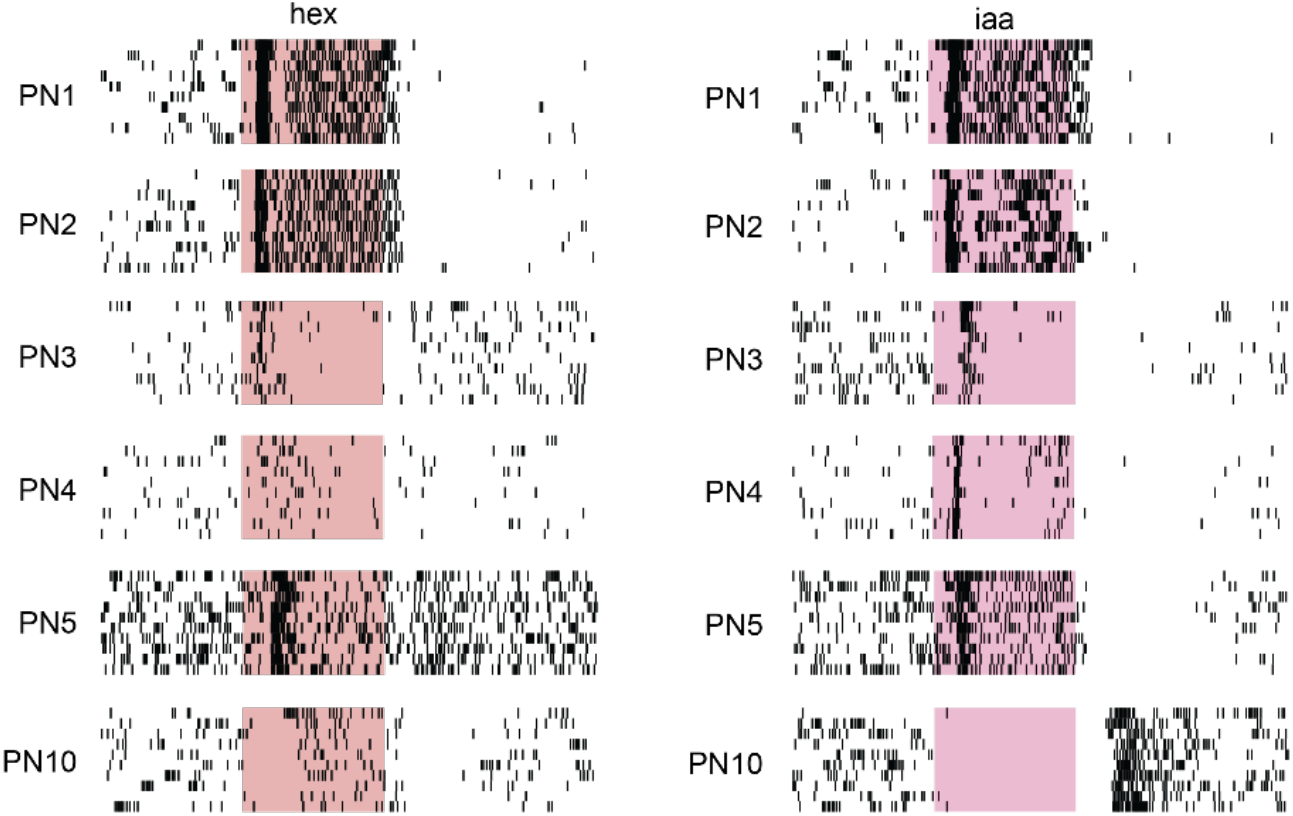
PN responses to 4 s solitary presentation of hexanol (hex; left) or isoamyl acetate (iaa; right) are shown. The colored boxes indicate the 4s duration of the odor pulse. Each tick corresponds to an action potential, and PN spiking activity is shown for 10 consecutive trials. Only PN responses to these solitary 4 s pulses was used to train all classification models (**Figures 3 – 5**) in the manuscript.

**Supplementary Figure 3:**
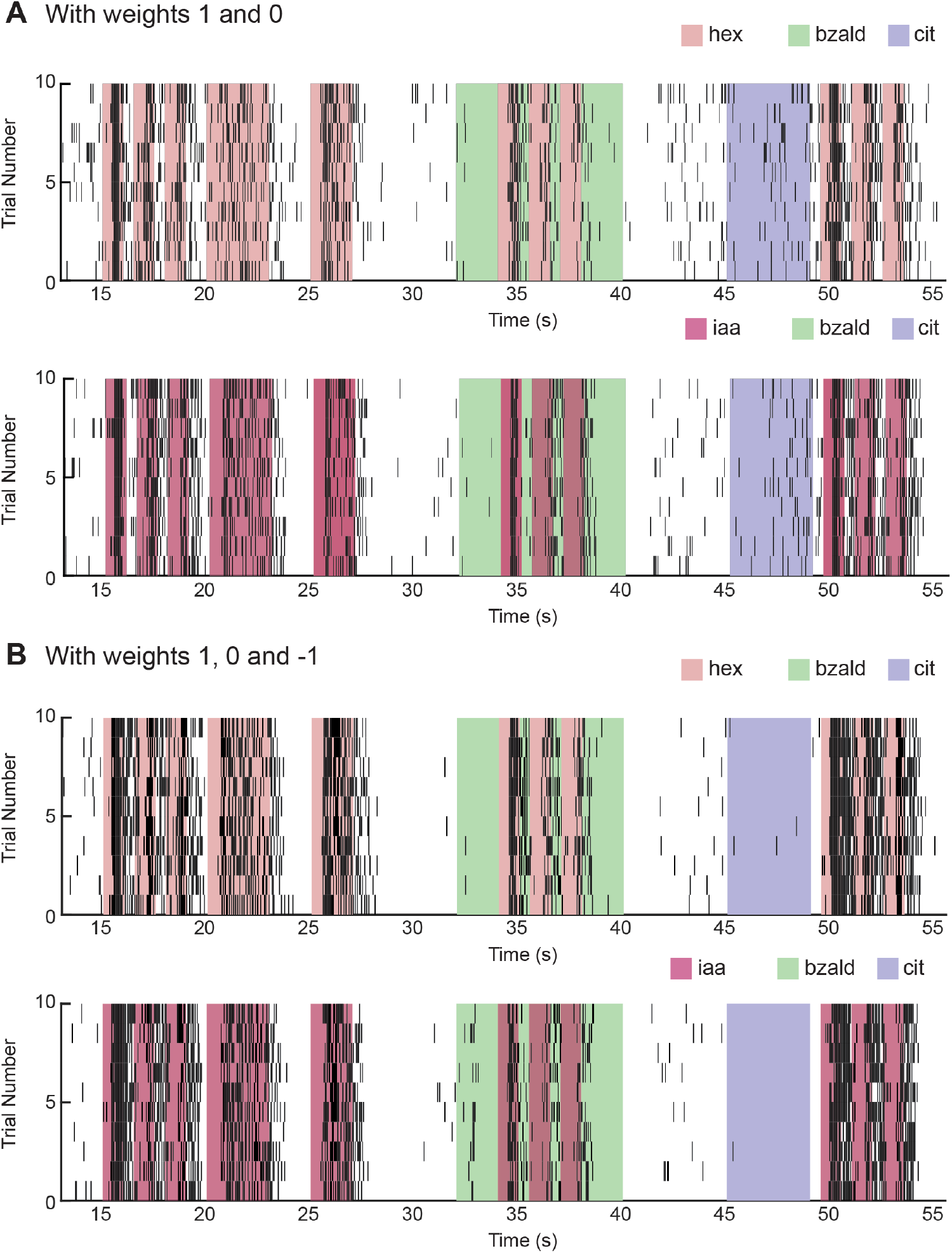
**(A)** Trial-by-trial prediction of hex presence using Bayesian logistic regression classifier with binary weights. Bottom panel shows similar results but for iaa classification problem. **(B)** Similar plot as in **panel A** but for Bayesian logistic regression approach with ternary weights {1 or 0 or -1}.

**Supplementary Figure 4:**
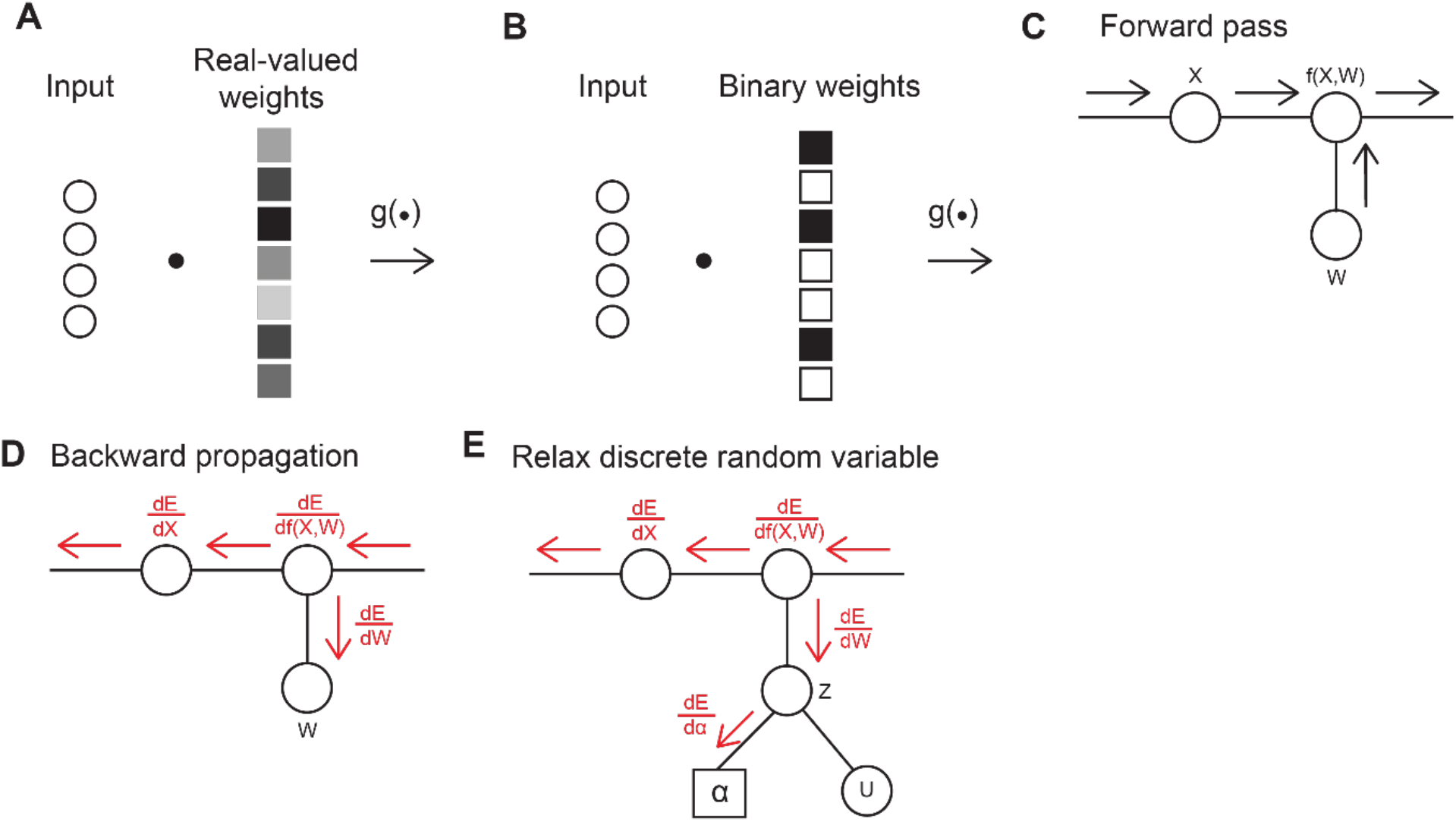
**(A)** Schematic of a regular feed-forward network model with just one layer. Network input or output from previous layer are weighted by real-valued weights and then the sum is sent to a non-linear activation function in the downstream nodes. **(B)** Schematic of a binary weighted network model. The network structure is same as before, but all the weights are constrained to be either 1 or -1. **(c)** The forward pass of neural network. All inputs traverse through all neurons layer-by-layer to the output of the neural network. **(D)** After the output and loss function is calculated. The gradient information is sent backward layer-by-layer to the first layer of neural network (i.e. backpropagation algorithm). Note that if one node is discrete, e.g. discrete weights, the gradient cannot be computed to update the weights. **(E)** Relax and reparameterize the discrete weights using fixed random variables and a parameterized function. The gradient information now can be used to update the parameters.

